# Development of a genetically encoded sensor for endogenous CaMKII activity

**DOI:** 10.1101/359398

**Authors:** Goli Ardestani, Megan West, Thomas J. Maresca, Rafael A. Fissore, Margaret M. Stratton

## Abstract

CaMKII is a crucial oligomeric enzyme in neuronal and cardiac signaling, fertilization and immunity. Here, we report the construction of a novel, substrate-based, genetically-encoded sensor for CaMKII activity, FRESCA (**FRE**T-based **S**ensor for **C**aMKII **A**ctivity). Currently, there is one biosensor for CaMKII activity, Camui, which contains CaMKII. FRESCA allows us to measure all endogenous CaMKII variants, while Camui can track a single variant. Since there are ~40 CaMKII variants, using FRESCA to measure aggregate activity allows a fresh perspective on CaMKII activity. We show, using live-cell imaging, FRESCA response is concurrent with Ca^2+^ rises in HEK293T cells and mouse eggs. In eggs, we stimulate oscillatory patterns of Ca^2+^ and observe the differential responses of FRESCA and Camui. Our results implicate an important role for the variable linker region in CaMKII, which tunes its activation. FRESCA will be a transformative tool for studies in neurons, cardiomyocytes and other CaMKII-containing cells.

## INTRODUCTION

Calcium-calmodulin dependent protein kinase II (CaMKII) is a serine/threonine kinase that plays critical signaling roles in multiple mammalian tissues (Backs et al., 2010; Rokita & Anderson, 2012; Shonesy, Jalan-Sakrikar, Cavener, & Colbran, 2014) and is implicated in a number of diseases (Mollova, Katus, & Backs, 2015; Robison, 2014; Steinkellner et al., 2012; Tu, Okamoto, Lipton, & Xu, 2014). CaMKII plays a key role in all electrically coupled cells, such as neurons and cardiomyocytes, and even cells that are not – such as lymphocytes and eggs – all of which communicate using Ca^2+^. Depending on the stimulus, the Ca^2+^ response leads is either a single Ca^2+^ rise or a more complex responses such as oscillations (Cuthbertson, Whittingham, & Cobbold, 1981; Eisner, Caldwell, Kistamas, & Trafford, 2017; Rutecki, 1992; Swann & Lai, 2013). Absence of Ca^2+^ signals causes severe defects in cell functionality, such as memory deficits in the case of neurons (Herring & Nicoll, 2016), or in the case of fertilization, failure to conceive (Escoffier et al., 2016; Yoon et al., 2008). CaMKII is responsible for reacting to Ca^2+^ oscillations and transducing this signal to downstream molecules. Indeed, it has been shown that neuronal CaMKII has a threshold frequency for activation (Chao et al., 2011; De Koninck & Schulman, 1998).

CaMKII has a unique oligomeric structure among the protein kinase family (Fig. 1A). Each subunit of CaMKII is comprised of a kinase domain, regulatory segment, variable linker region, and hub domain (Fig. 1B). The hub domain is responsible for oligomerization, which organizes into two stacked hexameric (or heptameric) rings to form a dodecameric (or tetradecameric) holoenzyme (Bhattacharyya et al., 2016; Chao et al., 2011; Rosenberg et al., 2006). In the absence of Ca^2+^, the regulatory segment binds to and blocks the substrate-binding pocket. Ca^2+^/calmodulin (Ca^2+^/CaM) turns CaMKII on by competitively binding the regulatory segment and exposing the substrate-binding pocket (Fig. 1C).

**Figure 1.**
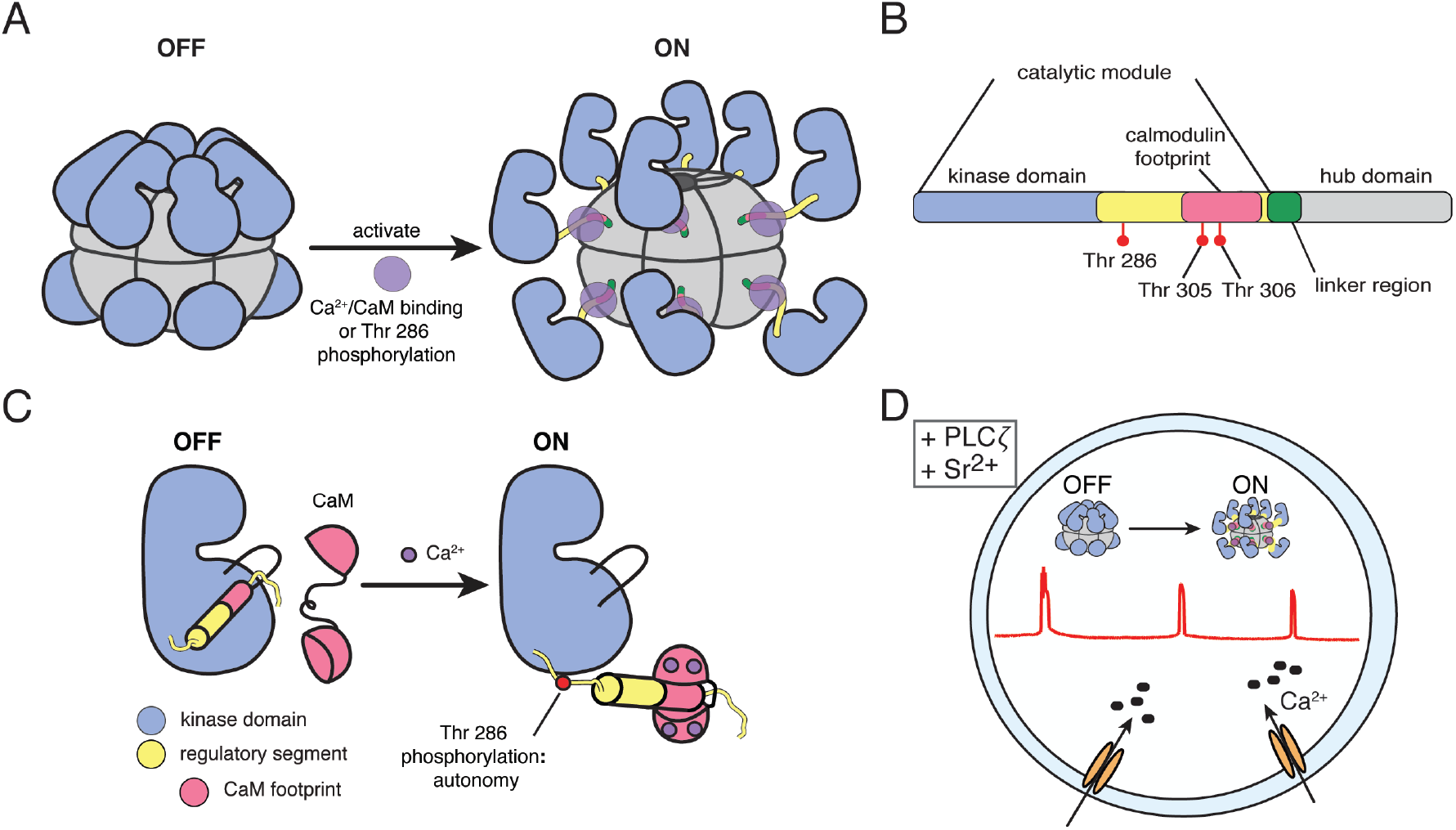
CaMKII: an essential enzyme. A) CaMKII is an oligomeric complex turned on by Ca^2+^/CaM binding, which facilitates phosphorylation at Thr 286. B) Each CaMKII subunit is comprised of a kinase domain, regulatory segment which houses the CaM binding domain, a variable linker region, and a hub domain. C) The regulatory segment of CaMKII maintains its off state in the absence of calcium by blocking its substrate binding pocket. Ca^2+^/CaM competes with the regulatory segment, thereby activating the kinase and allowing for Thr286 phosphorylation, which yields autonomous activity. Even when the calcium stimulus diminishes, CaMKII stays on as long as Thr 286 is phosphorylated. D) In mammalian eggs, addition of PLC*ζ* or Sr^2+^ leads to stimulation of calcium oscillations and CaMKII activation. CaMKII is expected to be “off” in the absence of Ca^2+^ and turn “on” after Ca^2+^ levels rise.

It has been demonstrated that CaMKII has a threshold frequency for activation (Chao et al., 2011; De Koninck & Schulman, 1998). There are four human CaMKII genes; CaMKIIα and β are predominantly expressed in neurons, CaMKIIδ is predominantly expressed in the heart and CaMKIIγ is found in multiple organ systems, including the reproductive organs. The kinase and hub domains of all four genes are highly conserved (~90% on average), however, the linker domain connecting the kinase and hub domains is variable in length and composition. Details elucidating the importance of the variable linker region remain to be uncovered, but there are >30 different splice variants of each of the four genes, which mostly vary in the linker region only. It has been shown that CaMKII activity is tuned by the length of the variable linker *in vitro* (Bayer, De Koninck, & Schulman, 2002; Chao et al., 2011). Specifically, as the variable linker is lengthened, less Ca^2+^ is needed for activation (*i.e.*, activation of CaMKII is easier). Thus, it is important for us to consider the complexity of endogenous CaMKII expressed in various cell types.

Camui is currently the only biosensor for CaMKII activity (Takao et al., 2005). Camui is a Förster resonance energy transfer (FRET)-based biosensor for CaMKII activity, which exploits the conformational change that CaMKII undergoes when it binds to Ca^2+^/CaM (Fig. 2A). To date, Camui has been a very useful tool to study and understand CaMKII activity in various cell types (mainly neurons and cardiomyocytes) and under various conditions (Erickson, Patel, Ferguson, Bossuyt, & Bers, 2011; Kwok et al., 2008; Takao et al., 2005). However, a major limitation is that the Camui sensor is constructed of a CaMKII variant itself, and thus will only report on this particular variant. To enhance our understanding of this complex protein, we need a way to measure endogenous CaMKII activity. One option is to re-engineer Camui with the appropriate CaMKII isoform to be studied, however this becomes limiting when there are multiple isoforms expressed in a single cell type, such as during the development of the female gamete, the egg. We now report the development of a novel biosensor that detects endogenous CaMKII activity. Herein, we show the efficacy of this new sensor in mouse eggs.

**Figure 2.**
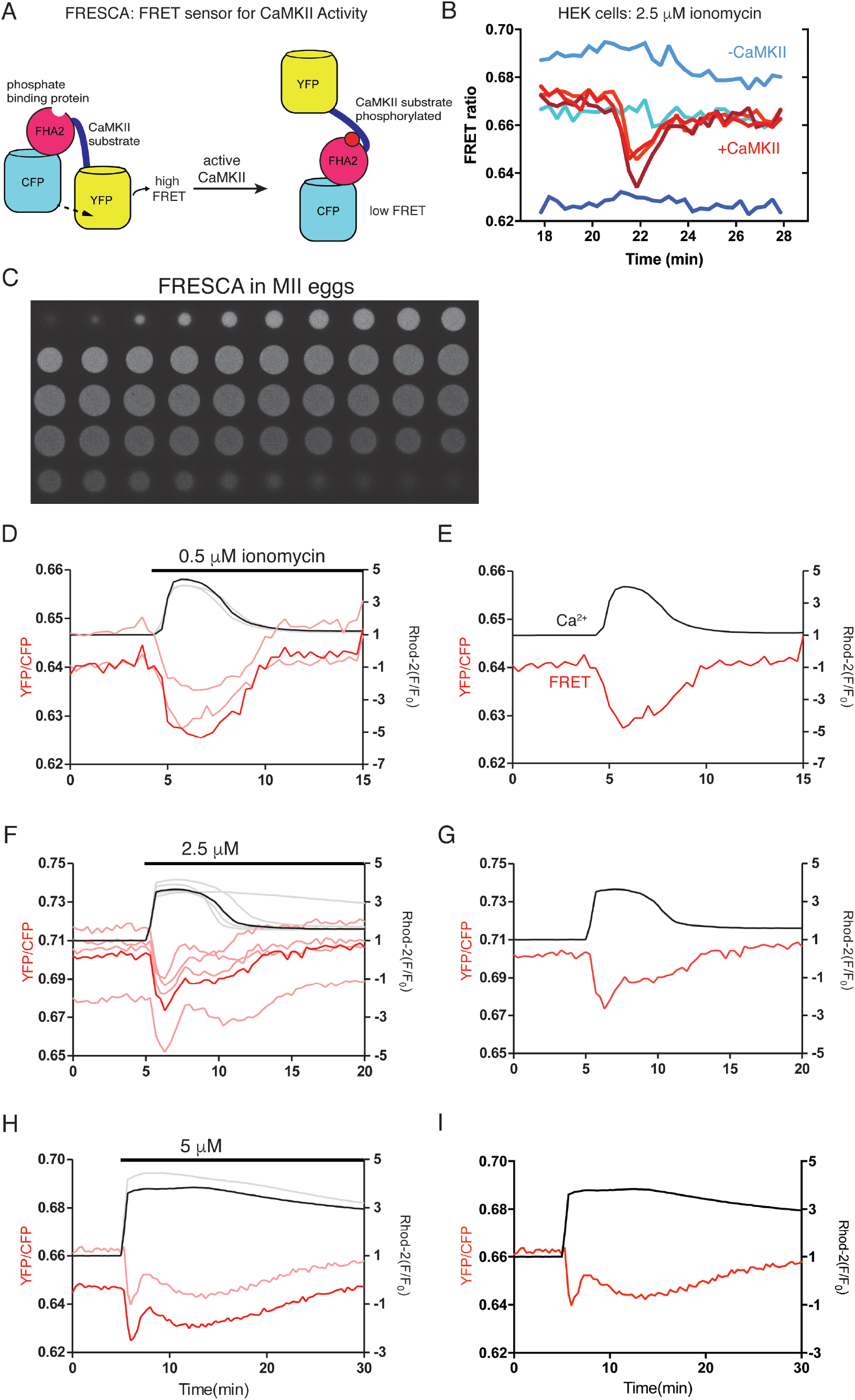
Monitoring endogenous CaMKII activity using FRESCA. A) A cartoon of a substrate-based CaMKII biosensor is shown: FRET Sensor for CaMKII Activity (FRESCA). Active CaMKII phosphorylates its substrate (syntide), which then acts as a substrate for FHA2 (phosphate binding domain). This induces a conformational change in the sensor as a consequence of CaMKII activity. B) FRESCA is transfected into HEK293 cells with and without CaMKII. Ionomycin (2.5 μM) is added to induce Ca^2+^ entry and FRET is monitored as YFP/CFP ratio. Without CaMKII transfected, there is no FRET change visualized. C) FRESCA expression in mouse MII eggs shows a widespread cytoplasmic distribution. D) Ca^2+^ is monitored using rhod-2 (black) and CaMKII activity is monitored using FRESCA (red). Multiple traces are shown after 0.5 μM ionomycin is added. E) One representative trace is shown after addition of 0.5 μM ionomycin. F) Multiple traces are shown after 2.5 μM ionomycin is added. G) One representative trace is shown after addition of 2.5 μM ionomycin. H) Multiple traces are shown after 5 μM ionomycin is added. I) One representative trace is shown after addition of 5 μM ionomycin.

## RESULTS AND DISCUSSION

### Development of a novel biosensor for endogenous CaMKII activity

We developed a novel substrate-based sensor for CaMKII activity, FRESCA (**<u>FRE</u>**T based **<u>S</u>**ensor for **<u>C</u>**aMKII **<u>A</u>**ctivity, Fig. 2A). We monitored CaMKII activity using FRESCA in real-time following the induction of Ca^2+^ responses to several agonists that are capable of initiating egg activation and embryogenesis. Building on a previous design for an Aurora kinase biosensor (Liu, Vader, Vromans, Lampson, & Lens, 2009), we replaced the sequence encoding the Aurora kinase substrate for the CaMKII substrate (syntide). The design also employs FHA2, a phosphate-binding domain, to facilitate a conformational change once the adjacent CaMKII substrate is phosphorylated (Durocher et al., 2000). FHA2 will bind to this phosphorylated Thr residue and produce a decrease in FRET between the terminal CFP/YFP pair.

#### Measuring the FRESCA response in HEK293T cells

We first tested the selectivity of FRESCA in HEK293T cells, which express negligible levels of CaMKII. We transfected HEK293T cells with either (i) CaMKII, calmodulin and FRESCA, or (ii) calmodulin and FRESCA. Ionomycin was added to the HEK293T cells to induce Ca^2+^ release and simultaneously monitored FRET (CFP/YFP ratio). We observed that with CaMKII present, the addition of ionomycin causes a reduction in FRET, indicating that CaMKII is active and phosphorylating FRESCA (Fig. 2B, red lines). Importantly, we did not observe a FRET change when CaMKII was not co-transfected, demonstrating that FRESCA is selective for CaMKII and not being phosphorylated by other HEK cell kinases (Fig. 2B, blue lines).

#### Using FRESCA to monitor CaMKII activity in mouse eggs

HEK293T cells provided a good model for a highly controlled evaluation of CaMKII activity and the ability of FRESCA to specifically report on CaMKII in the presence of other cellular kinases. However, we wanted to test FRESCA in a more complex and native system, importantly, with endogenous CaMKII and where the Ca^2+^ response has a clear physiological function. To accomplish this, we expressed FRESCA in mouse eggs to measure endogenous CaMKIIγ activity following increases in intracellular Ca^2+^ induced by a variety of agonists.

In all mammals, Ca^2+^ oscillations are required for initiation of embryogenesis (Deguchi, Shirakawa, Oda, Mohri, & Miyazaki, 2000; Fissore, Dobrinsky, Balise, Duby, & Robl, 1992). Mammalian eggs are arrested at metaphase II of meiosis; both Ca^2+^ oscillations and consequent CaMKII activity are required for release from this arrest (Fig. 1D) (Backs et al., 2010; Chang, Minahan, Merriman, & Jones, 2009; Miao, Stein, Jefferson, Padilla-Banks, & Williams, 2012; Miyazaki et al., 1992; Presler et al., 2017). Ca^2+^ oscillations are induced following gamete fusion when the sperm releases into the egg a sperm specific protein (PLCzeta; *ζ*), which triggers the Ca^2+^ responses (Ducibella et al., 2002; Saunders et al., 2002). On average, there is one Ca^2+^ rise every 20 minutes and oscillations in mouse zygotes last for ~4 hours, which coincides with the formation of the pronuclei (PN) (Jones, Carroll, Merriman, Whittingham, & Kono, 1995). The source of this Ca^2+^ is from internal stores, which are replenished by Ca^2+^ influx from the extracellular media. CaMKII is activated simultaneously with the initiation of Ca^2+^ oscillations and female mice that are CaMKIIγ null are sterile (Backs et al., 2009). Despite the role of CaMKIIγ in the initiation of development, the complete profile of CaMKII activity during fertilization in mammals is not known. Further, CaMKII activity also seems to play a role in preventing apoptosis in *Xenopus* and mouse eggs, although the pattern and degree of activation for this activity are even less studied (Nutt et al., 2005).

To date, CaMKII activity has only been assessed based on a few Ca^2+^ rises using *in vitro* kinase assays and during only the first hour of oscillations, which is considerably shorter than the time scale for normal oscillations in the mouse. Therefore, there is a need to monitor CaMKII activity in live cells and for an extended time, which is what we address here.

#### FRESCA and ionomycin-induced Ca^2+^ oscillations in mouse eggs

FRESCA expression and distribution in germinal vesicle (GV) oocytes and MII stage oocytes, henceforth referred to as eggs, was widespread and cytoplasmic (Fig. 2C). However, a small amount of FRESCA appeared to enter the nucleus of GV oocytes (Fig. 2, supplement 1).

Given the immediate and large Ca^2+^ rise caused by the addition of ionomycin, we first tested FRESCA responses in eggs using this ionophore. We analyzed the effect of 3 concentrations of ionomycin: 0.5 μM, 2.5 μM and 5 μM. Upon addition of ionomycin to eggs expressing FRESCA, we observed a FRET decrease, indicating CaMKII activity (Fig. 2D-I). At the lowest ionomycin concentration (0.5 μM), CaMKII activity appears to perfectly track the Ca^2+^ pulse (Fig. 2D, E). Conversely, at higher ionomycin concentrations, CaMKII activity is unstable during the duration of the Ca^2+^ pulse, although higher concentrations appeared to prolong and increase the FRET response of FRESCA (Fig. 2, supplement 2). The time to FRET peak was faster with addition of higher ionomycin concentrations (Fig. 2, supplement 2).

We tested the specificity of FRESCA for CaMKII in mouse eggs. We first used CaMKII inhibitors, which should eliminate the FRET response if CaMKII is the only kinase phosphorylating FRESCA in eggs (Fig. 2, supplement 3). We show that the addition of KN93, a commonly used allosteric inhibitor for CaMKII, significantly reduces FRET (Madgwick, Levasseur, & Jones, 2005; Smyth et al., 2002). Addition of AS105, an ATP competitive CaMKII specific inhibitor, also significantly reduces FRET, while its inactive analog (AS461) does not affect FRET (Neef et al., 2018). We also tested inhibition and activation of protein kinase C (PKC), since PKC is the other major Ca^2+^ sensitive kinase in mouse eggs (Medvedev, Stein, & Schultz, 2014; Wang et al., 2010). Addition of PKC inhibitors Bim1 (Halet, 2004) and GO6983 (Gou, Wang, Zou, Qi, & Xu, 2018) do not affect the FRESCA signal, indicating that PKC is not phosphorylating FRESCA. Eggs do not express conventional PKC isoforms, so the results of the broad-spectrum PKC inhibitor (GO6983) reinforced the results of Bim1. Finally, addition of PMA, a PKC activator shown to stimulate this enzyme in mouse eggs (Halet, 2004), also does not induce FRET (Fig. 2, supplement 3). Taken together, we report that FRESCA is a specific reporter of CaMKII activity in mouse eggs.

#### FRESCA and Sr^2+^-induced Ca^2+^ oscillations

Addition of 10 mM Sr^2+^ to the extracellular media in place of external Ca^2+^ is a common method of parthenogenetic activation in mouse eggs, and it induces highly consistent oscillations in these cells (Fig. 3, black lines); these oscillations initiate all events of egg activation (Bosmikich & Whittingham, 1995; Carvacho, Lee, Fissore, & Clapham, 2013; Kline & Kline, 1992). Further, the TRPV3 channel has recently been identified as the channel responsible for Sr^2+^ influx in mouse eggs (Carvacho et al., 2013). We therefore examined the response of endogenous CaMKII to Sr^2+^-induced oscillations. We observed endogenous CaMKIIγ activity (monitored by FRESCA) almost simultaneously with the initiation of oscillations. Indeed, nearly all eggs (11/13) showed CaMKII activity within the first two rises. Importantly, CaMKII activity is reproduced over time, as FRESCA continues to track each Ca^2+^ rise for >2 hours (Fig. 3B).

**Figure 3.**
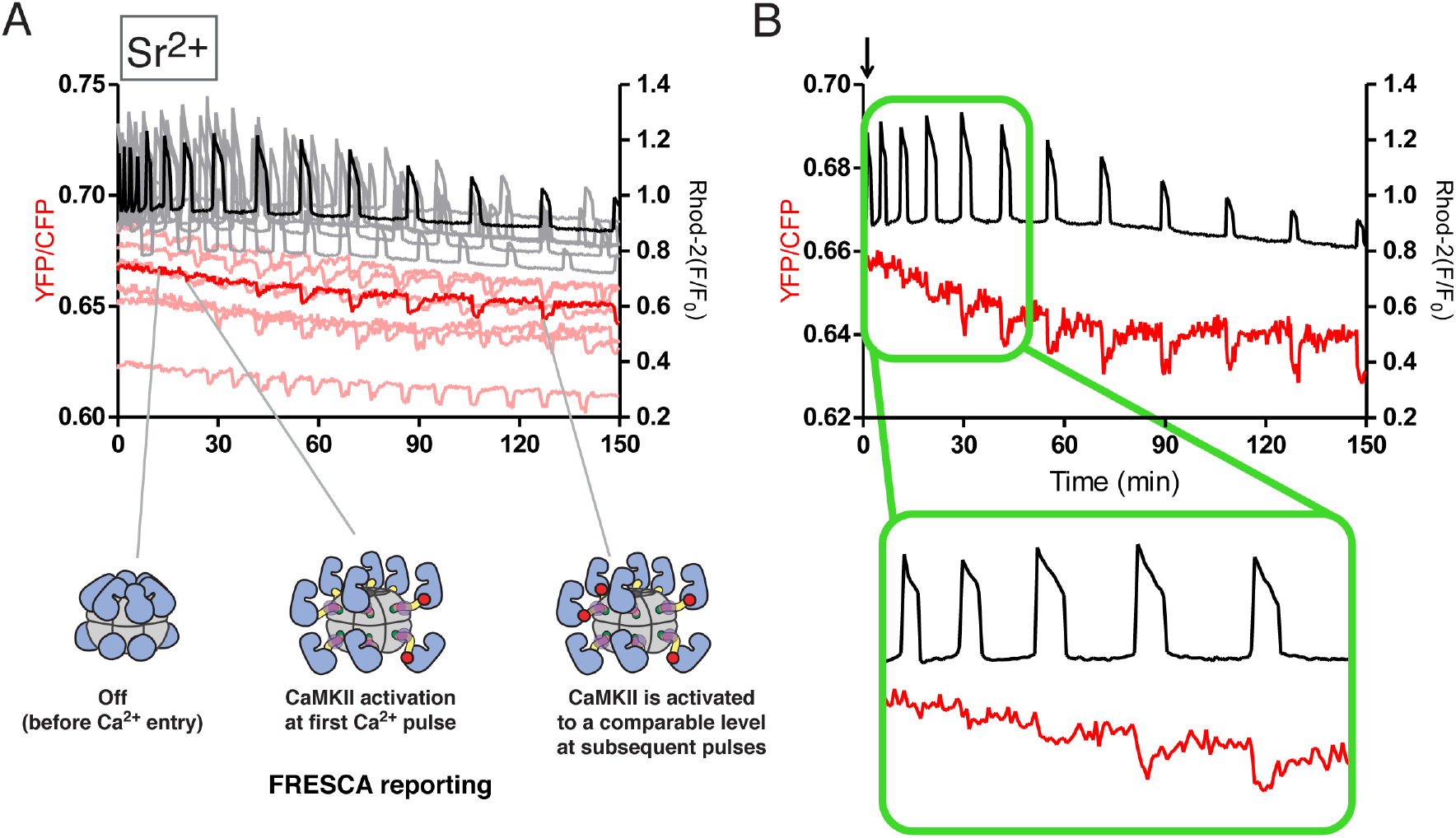
Endogenous CaMKII activity tracks Ca^2+^ oscillations in mouse eggs. A) Ca^2+^ oscillations in eggs are induced by addition of Sr^2+^ to the extracellular media. Ca^2+^ is monitored by Rhod-2 (black line) and endogenous CaMKII activity is tracked by FRESCA (red line). B) One representative trace from Sr^2+^ oscillations is shown. Inset focuses on the first Ca^2+^ rises.

We propose a molecular model to describe this data. The Ca^2+^ rises progressively decrease in duration/amplitude over time (Deguchi et al., 2000), however, the FRESCA responses are not diminished and the peak kinase activity seems to outlast the peak elevation of Ca^2+^/Sr^2+^. This suggests autophosphorylation of CaMKII at Thr 286, which facilitates activation at subsequent Ca^2+^ pulses by increasing the affinity for Ca^2+^/CaM (see cartoons in Fig. 3A) (Meyer, Hanson, Stryer, & Schulman, 1992). Additionally, a prolonged time course of FRESCA response to Sr^2+^ indicates that FRESCA continues to faithfully track endogenous CaMKII up to 6 hours (Fig. 3, supplement 2).

### Using Camui to measure CaMKII activity in mouse eggs

As described, the highly used Camui biosensor is comprised of CaMKII itself (see Fig. 4A), specifically CaMKIIα, which has a 30-residue variable linker region (Fig. 4C). Despite its widespread use, Camui has not yet been used to monitor CaMKII activity in mouse eggs. Given that it has been demonstrated that CaMKII activity is tuned by the length of the variable linker (Bayer et al., 2002; Chao et al., 2011), specifically, as the variable linker is lengthened, less Ca^2+^ is needed for activation, we hypothesized that FRESCA may report CaMKII activity in mouse eggs more faithfully than Camui. This assumption is based on the knowledge that mouse eggs express equimolar concentrations of the two versions of CaMKIIγ (γ3 and γJ), which have 69 and 90 residue variable linkers, respectively (Fig. 4C) (Hatch & Capco, 2001; Suzuki, Hara, Takagi, Yamamoto, & Ueno, 2011), considerably longer than the 30-residue linker of CaMKIIα. We therefore tested the response of Camui compared to FRESCA.

**Figure 4.**
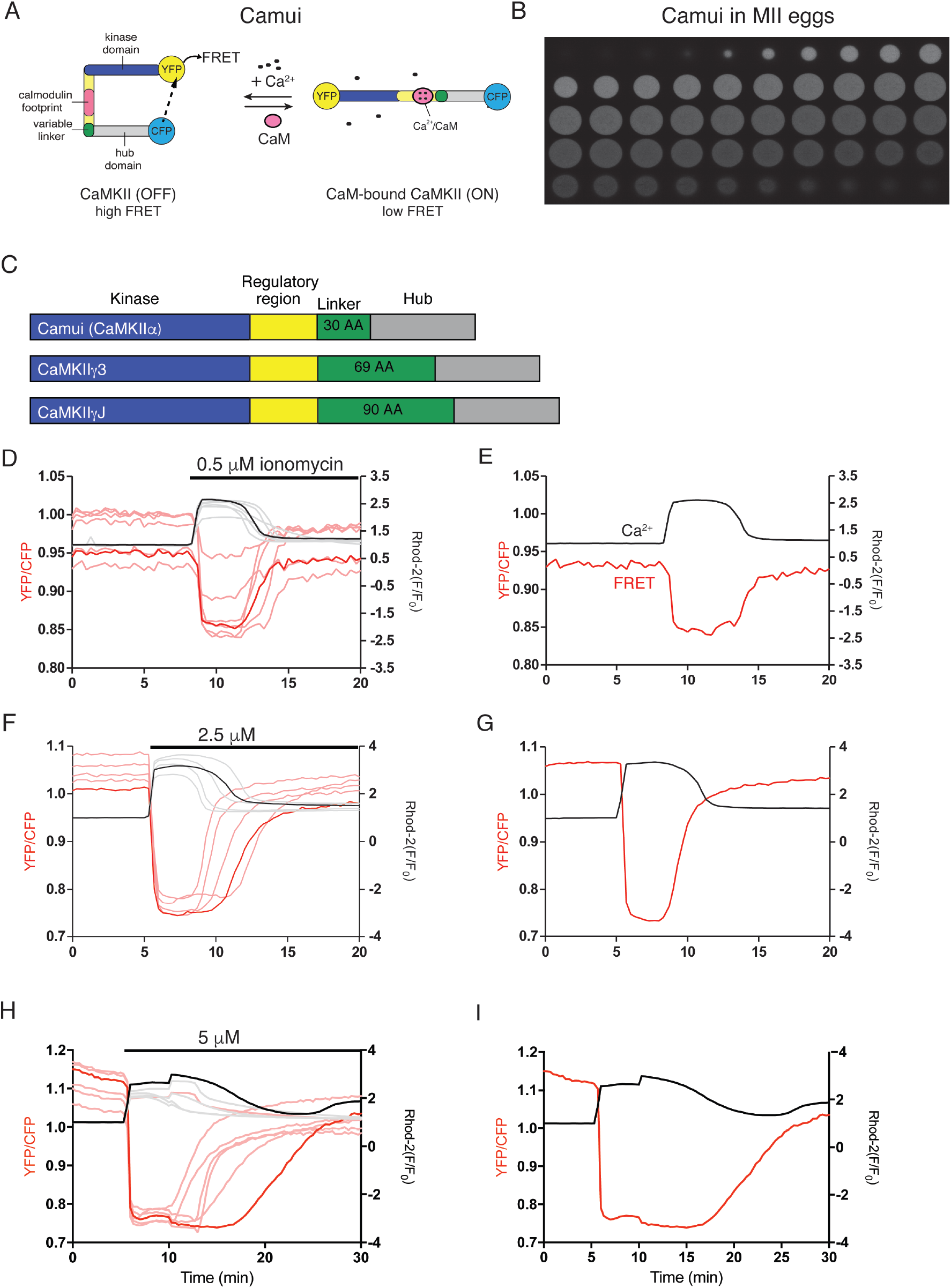
Simultaneously monitoring Ca^2+^ influx and CaMKII activity using Camui. A) Camui is an existing biosensor for CaMKII activity, which exploits the conformational change of CaMKII binding to Ca^2+^/CaM to report on activity using FRET. B) Camui expression in mouse MII eggs shows a widespread cytoplasmic distribution. C) Linear sequences of Camui (CaMKIIα), and the two CaMKII isoforms reported in eggs (CaMKII γ3 and γJ). D) Changes in Ca^2+^ are monitored using rhod-2 (black) and CaMKII activity is monitored using Camui (red). Multiple traces are shown after 0.5 μM ionomycin is added. E) One representative trace is shown after addition of 0.5 μM ionomycin. F) Multiple traces are shown after 2.5 μM ionomycin is added. G) One representative trace is shown after addition of 2.5 μM ionomycin. H) Multiple traces are shown after 5 μM ionomycin is added. I) One representative trace is shown after addition of 5 μM ionomycin.

We expressed the Camui reporter in mouse eggs using mRNA injection, and similar to FRESCA, expression was robust within ~30 minutes and we began FRET measurements ~4 hours post injection to attain stable Camui levels. As shown by confocal microscopy, Camui attained a widespread cytoplasmic expression in eggs, although in GVs it was excluded from the nucleus, which is consistent with the reported expression of CaMKII in the cytosol of mouse eggs (Fig. 4B and Fig 4., supplement 1) (Hatch & Capco, 2001).

#### Camui and ionomycin-induced Ca^2+^ oscillations

We first induced Ca^2+^ release in eggs by adding ionomycin as previously described and simultaneously monitored changes in FRET values (YFP/CFP) (Fig. 4C-H). As with FRESCA, a decrease in FRET is indicative of an increase in CaMKII activity. It is clear that CaMKII activity increases (red line) coincident with the increase in Ca^2+^ (black line) in all conditions. The Ca^2+^ and Camui responses increased dose-dependently and approximately synchronously, as the large increase in the amount of Ca^2+^ release caused by increasing ionomycin from 0.5 μM to 2.5 μM, results in a 1.9-fold increase in CaMKII activity (mean amplitude of FRET change) (Fig. 4, supplement 2). Further increasing ionomycin from 2.5 μM to 5 μM produces nearly no change in total Ca^2+^ release, although it is very likely that the reporting range of Rhod-2 is saturated at these levels of Ca^2+^ release. The CaMKII activity also appears to remain constant, although this may also represent saturation of the FRET signal (Fig. 4, supplement 2). Notably, addition of 5 μM ionomycin results in a prolonged duration of activity compared to lower concentrations, but it is unclear whether this reflects the extended activation of the enzyme or cellular stress.

The absolute amplitude of the change in the FRET ratio for Camui after the addition of 0.5 μM ionomycin is ~10-fold greater than what is observed for FRESCA (0.014 for FRESCA compared to 0.14 for Camui), however, this measurable signal change in FRESCA is sufficient to monitor endogenous CaMKIIγ activity, which we cannot detect with Camui, which reports CaMKIIα activity. In addition, it is worth noting that the shape of the FRESCA traces is slightly different from those of Camui for the same stimulus. The peak activity of FRESCA is shorter than the corresponding Ca^2+^ peak, although the return to basal activity is more protracted, whereas the Camui response more perfectly tracks the shape of the Ca^2+^ peak.

#### Camui and Sr^2+^ induced Ca^2+^ oscillations

We next examined the Camui response to Sr^2+^-induced oscillations (Fig. 5). Remarkably, despite the presence of robust changes in intracellular Ca^2+^ levels, Camui (Fig. 5, red line) did not report any CaMKII activity until the ~6^th^ significant Ca^2+^ rise (Fig. 5B, arrow and inset). In total, only 33% of the eggs expressing Camui showed activity in the first two rises, whereas nearly all FRESCA expressing eggs showed activity within the first two rises. These data suggest that the endogenous CaMKIIγ in eggs is more sensitive to Ca^2+^/CaM than CaMKIIα. This finding is in line with previous data showing that longer linker CaMKII splice variants (CaMKIIγ3 and CaMKIIγJ) are activated at lower concentrations of Ca^2+^/CaM than shorter linker variants (CaMKIIα) (see Fig. 4C) (Chao et al., 2011). Additionally, the delayed response seen in the Camui eggs could also be a result of endogenous CaMKII being activated first (lower EC_50_ for Ca^2+^/CaM) thereby competing with Camui for the available activating ligand.

**Figure 5.**
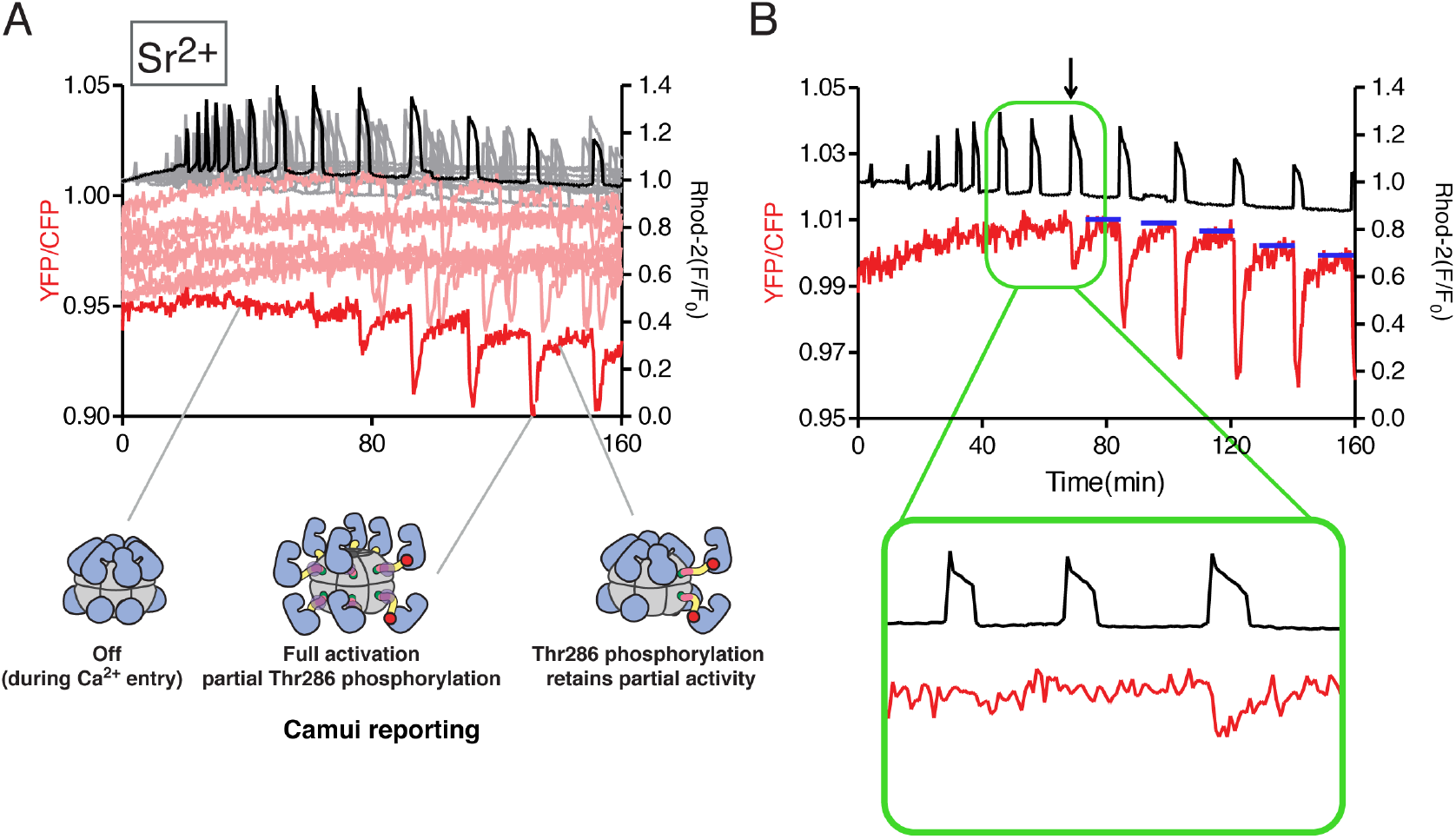
Camui activity tracks Ca^2+^ oscillations in mouse eggs in a delayed manner. A) Ca^2+^ oscillations are induced by addition of Sr^2+^. Ca^2+^ is monitored by Rhod-2 (black line) and CaMKII activity is tracked by Camui (red line). CaMKII cartoons indicate hypothesized molecular details during the Ca^2+^ pulses. B) One representative trace from Sr^2+^ oscillations is shown. Blue lines indicate baseline CaMKII activity after each rise. Arrow indicates first significant FRET response.

Another distinctive feature of the Camui response caused by Sr^2+^ oscillations is that whereas the initial Camui responses were delayed, once they commenced, they displayed an integrated activation with each subsequent pulse. For example, we analyzed the mean amplitude for the first three observable FRET changes. From the first to the second FRET change, there was a 1.6-fold increase in CaMKII activity. From the second to the third FRET change, there was a negligible change, and these changed occurred while the amplitude of the Ca^2+^ peaks progressively decreased and/or remained unchanged (Fig. 5, supplement 1). These data indicate that CaMKII activity, once stimulated, is cooperative with each additional Ca^2+^ pulse. This result is consistent with previous data showing that CaMKII activity is highly cooperative *in vitro* (Chao et al., 2010; Chao et al., 2011). As depicted in Figure 5A, a potential explanation for this is phosphorylation at Thr286 which may persist even in the absence of elevated Ca^2+^. It has been clearly shown that CaMKII with Thr286 phosphorylated has a significantly higher affinity for Ca^2+^/CaM (Meyer et al., 1992). This would also explain why the FRET level does not return to baseline in between later Ca^2+^ oscillations (Fig. 5B, blue lines).

### Measuring CaMKII activity under native fertilization conditions

In mammals, fertilization-associated Ca^2+^ oscillations are induced by the release of sperm’s PLC*ζ* into the ooplasm (Saunders et al., 2002). We tested the response of both FRESCA and Camui in response to the expression of PLC*ζ*.

#### FRESCA and PLC*ζ*-induced Ca^2+^ oscillations

We accomplished PLC*ζ* expression by injection of its mRNA into FRESCA expressing eggs, and thereafter began monitoring changes in FRESCA responses (Fig. 6A, B). The initiation of oscillations stimulated the early activity of the endogenous CaMKIIγ, and this activity was detected with each additional rise. Similar to Sr^2+^ induced oscillations, we observed a relative decrease in the amplitude of the Ca^2+^ pulses over time, yet the FRESCA response was largely maintained (Fig. 6, supplement 1). These observations are the longest evaluation of CaMKII activity following natural oscillations ever reported following fertilization, as previous studies only reported up to 60 min post-initiation of oscillations (Markoulaki, Matson, & Ducibella, 2004; Ozil et al., 2005).

**Figure 6.**
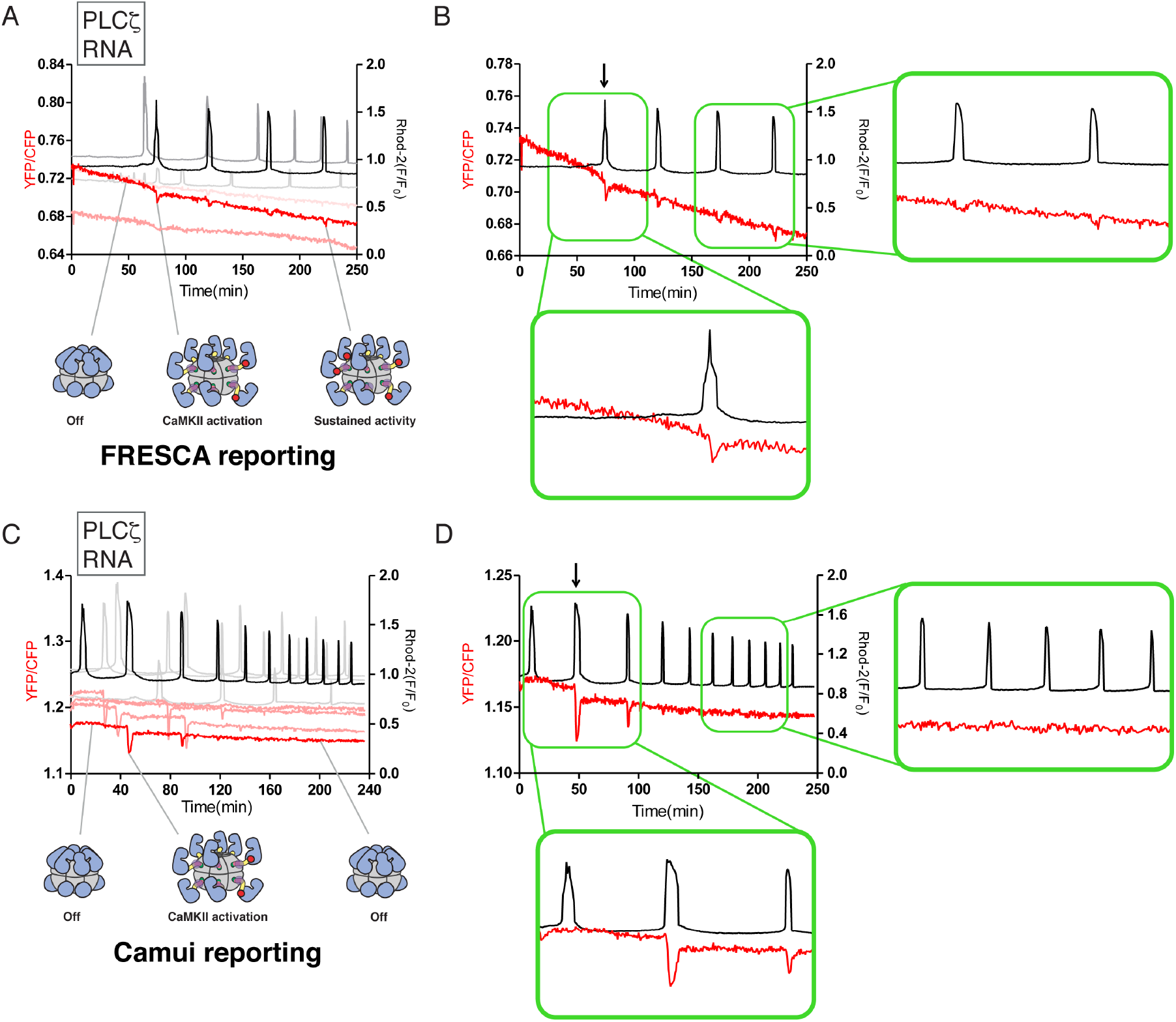
FRESCA, but not Camui, continues to report CaMKII activation by Ca^2+^ oscillations induced by PLC*ζ* injection. Ca^2+^ oscillations are induced by injection of PLC*ζ* mRNA. CaMKII activity is monitored using FRESCA or Camui (FRET, red lines) and Ca^2+^ is monitored using Rhod-2 (black lines). A) An overlay of 3 representative eggs using FRESCA as the reporter of endogenous CaMKII activity. Cartoon depictions of hypothesized states of CaMKII are shown below. Red circles indicate Thr286 phosphorylation. B) One representative trace from PLC*ζ*-induced oscillations and FRESCA reporting is shown. Insets highlight the first and last pulses. C) An overlay of 4 representative eggs using Camui as the reporter of CaMKIIα activity. Cartoon depictions of hypothesized states of CaMKII are shown below. Red circles indicate Thr286 phosphorylation. D) One representative trace from PLC*ζ*-induced oscillations and Camui reporting is shown. Insets highlight the first and last pulses.

#### Camui and PLC*ζ*-induced Ca^2+^ oscillations

We next assessed how Camui would report CaMKII activity induced by Ca^2+^ oscillations, and compare the response to those induced by Sr^2+^. To do this, eggs expressing Camui were injected with PLC*ζ* mRNA and Ca^2+^ and FRET responses were monitored. Similar to FRESCA, we observed that Ca^2+^ oscillations nearly immediately induced CaMKII activity as monitored by Camui (Fig. 6D, arrow and bottom inset). However, this initial activity was not detected in subsequent rises, and only the first and second (and to a less extent, third) Ca^2+^ rises induced Camui responses despite the presence of robust and frequent Ca^2+^oscillations (Fig. 6C, D, supplement 2). Additionally, it is worth pointing out that the area under the curve for the third Ca^2+^ rise in these experiments was significantly reduced. This may be due to the fact that Camui itself is contributing significantly to the existing CaMKII in the egg, and potentially altering Ca^2+^ dynamics. These results raised the possibility that Camui is not well suited to detect CaMKII activity initiated by sporadic and low magnitude Ca^2+^ rises, which are characteristic of mammalian fertilization. Regardless, it remains to be elucidated why Sr^2+^ induced oscillations are able to protractedly promote robust and persistent Camui responses whereas the Camui response to PLC*ζ*-induced oscillations fades rapidly.

## CONCLUDING REMARKS

It has been appreciated for decades that both Ca^2+^ oscillations and CaMKII activation in mouse eggs is crucial to fertilization and initiation of embryo development. Here, we provide an analysis of CaMKII activation in real-time in eggs using FRET-based CaMKII biosensors. Importantly, our new biosensor, FRESCA, allowed us to monitor endogenous CaMKII (CaMKIIγ3 and γJ) activation in real-time as a consequence of different activation stimuli (ionomycin, Sr^2+^, and PLC*ζ*). The FRESCA response was noticeably different from the Camui sensor, which reports on CaMKIIα. When different Ca^2+^ oscillation patterns are induced, we observe subsequent differences in CaMKII activity. From our data, it is clear that both the (i) pattern of Ca^2+^ oscillations as well as the (ii) specific CaMKII isoform responding play a role in CaMKII activation.

The pivotal role of CaMKII activation in causing release of the meiotic arrest and activation of the embryonic developmental program in vertebrates was recently and more specifically evidenced by careful mass spectrometry experiments (Presler et al., 2017). This study showed that soon after fertilization, and temporally coinciding with the Ca^2+^ wave, there is a strong increase in protein phosphorylation that far outweighs the biochemical changes caused by protein degradation that accompanies fertilization. Remarkably, the study also found that 25% of the phosphorylated sites matched the minimal phosphorylation motif of CaMKII. It is therefore important to determine how Ca^2+^ rises turn on CaMKII activity, and what parameter(s) of individual rises within an oscillatory pattern are necessary for periodic and consistent stimulation of its activity. We propose that the magnitude of the initial activation of CaMKII depends on the magnitude of the stimulus and on internal regulation of CaMKII, which is largely based on the variable linker region. Knowing the minimal Ca^2+^ signal that increases the activity of CaMKIIγ is important as we seek to develop more physiological methods of parthenogenetic activation to treat some cases of infertility.

More broadly, now that we have demonstrated the utility of FRESCA in mouse eggs, this opens the door to measuring endogenous CaMKII activity in other cell types, such as neurons and cardiomyocytes. CaMKII activation has been heavily studied *in vitro* (Chao et al., 2010; Chao et al., 2011; Rosenberg, Deindl, Sung, Nairn, & Kuriyan, 2005), and it is intriguing to also consider the potential effects of subunit exchange in cellular conditions (Bhattacharyya et al., 2016; Stratton et al., 2013). It will be necessary to increase the signal to noise ratio of the FRESCA sensor in order to achieve a more robust signal for accurate quantification of kinetics and amplitudes. Once this is accomplished, we believe that FRESCA will provide new insights into CaMKII activity in cells and allow us to unravel the complexity of this unique protein kinase.

## Acknowledgements

We thank Changli He for assistance on RNA purification. We thank Allosteros therapeutics for providing AS105, AS461 and Howard Schulman for helpful discussion. We also thank Peter Chien, Eric Strieter, Scott Garman for discussions and John Kuriyan for helpful comments on the manuscript.

## MATERIALS AND METHODS

### Plasmid design

In order to accommodate the requirements for FHA2 binding (Durocher et al., 2000), syntide was modified from PLARTLSVAGLPGKK to PLARALTVAGLPGKK to create syntide-2. Syntide-2 was generated by annealing GATCCGGCGGCGCCGGCGGCGGCccgctggcgcgcgccctgaccgtggcgggcctgccgggcaaaaa aGGC and GGCCGCCttttttgcccggcaggcccgccacggtcagggcgcgcgccagcggGCCGCCGCCGGCGCCG CCG (IDT), which produced *BamHI* site at the 5’ end and a *NotI* site on the 3’ end. This product was phosphorylated (Ambion Pnk), purified (Thermo Fisher) and then ligated using T4 DNA ligase (Invitrogen) into a plasmid encoding the Aurora kinase FRET sensor (kind gift from Thomas Maresca). The final FRESCA sensor (with syntide-2 in place of the Aurora substrate) was cloned into pCDNA3.1.

### HEK 293T Cell culture

All HEK293T cell cultures were grown in Dulbecco’s Modified Eagle’s Medium (Sigma) supplemented with 10% fetal bovine serum (Sigma) and maintained at 37°C and 5% carbon dioxide levels. The identity of these cells was authenticated by ATCC (CRL-3216 ATC 293T, Lot #63226319). Cells were transfected using Lipofectamine^®^ 2000 Reagent (Invitrogen) and 150 ng of DNA constructs.

### Collection of mouse eggs

Metaphase II (MII) eggs were collected from the oviducts of 6- to 10-week-old CD-1 female mice 12–14 h after administration of 5 IU of human chorionic gonadotropin (hCG), which was administered 46–48h after the injection of 5 IU of pregnant mare serum gonadotropin (PMSG; Sigma; Saint Louis, MO). Cumulus cells were removed with 0.1% bovine testes hyaluronidase (Sigma). MII eggs were placed in KSOM with amino acids (Millipore Sigma) under mineral oil at 37°C in a humidified atmosphere of 5% CO_2_ until the time of monitoring. All animal procedures were performed according to research animal protocols approved by the University of Massachusetts Institutional Animal Care and Use Committee.

### Preparation of cRNAs and Microinjections

The sequences encoding Camui and FRESCA were subcloned into a pcDNA6 vector (pcDNA6/Myc-His B; Invitrogen, Carlsbad, CA) between the XhoI and PmeI restriction sites. Mouse PLC*ζ* was a kind gift from Dr K. Fukami (Tokyo University of Pharmacy and Life Science, Japan) and subcloned into a PCS2+ vector, as previously described by us (Kurokawa et al., 2007). Plasmids were linearized with a restriction enzyme downstream of the insert to be transcribed and cDNAs were *in vitro* transcribed using the T7 or SP6 mMESSAGE mMACHINE Kit (Ambion, Austin, TX) according to the promoter present in the construct. A Poly (A)-tail was added to the mRNAs using a Tailing Kit (Ambion) and poly(A)-tailed RNAs were eluted with RNAase-free water and stored in aliquots at −80 °C. Microinjections were performed as described previously (Lee *et al.,* 2016). cRNAs were centrifuged, and the top 1–2 μl was used to prepare micro drops from which glass micropipettes were loaded by aspiration. cRNA were delivered into eggs by pneumatic pressure (PLI-100 picoinjector, Harvard Apparatus, Cambridge, MA). Each egg received 5–10 pl, which is approximately 1–3% of the total volume of the egg. Injected MII eggs were allowed for translation up to 4h in KSOM. Group of eggs were injected with mouse PLC*ζ* after 4h of FRET construct injection.

### FRET and Calcium imaging

To estimate relative changes in the cytoplasmic activity of Camui and/or FRESCA, emission ratio imaging of the Camui and FRESCA (YFP/CFP) was performed using a CFP excitation filter, dichroic beam splitter, CFP and YFP emission filters (Chroma technology, Rockingham, VT; ET436/20X, 89007bs, ET480/40m and ET535/30m). To measure Camui and/or FRESCA activity and [Ca^2+^]_i_ simultaneously, eggs that had been injected with Camui and/or FRESCA cRNAs were loaded ~ 4 hours post-injection with 1 μM Rhod-2AM supplemented with 0.02% pluronic acid for 20 minutes at RT. Eggs were then attached on glass-bottom dishes (MatTek Corp., Ashland, MA) and placed on the stage of an inverted microscope. CFP, YFP and Rhod-2 intensities were collected every 20 second by a cooled Photometrics SenSys CCD camera (Roper Scientific, Tucson, AZ). The rotation of excitation and emission filter wheels was controlled using the MAC5000 filter wheel/shutter control box (Ludl) and NIS-elements software (Nikon). Imaging was performed on an inverted epifluorescence microscope (Nikon Eclipse TE 300, Analis Ghent, Belgium) using a 20x objective. For studies where ionomycin was used to induce Ca^2+^ responses, eggs were transferred into a 360 μl Ca^2+^-free TL-Hepes drop on a glass bottom dish, after which and following a brief monitoring period to determine baseline [Ca^2+^]_i_ values, different concentrations of ionomycin were added and Ca^2+^ responses monitored. For Sr^2+^ studies, eggs were transferred into a Ca^2+^-free TL-Hepes, containing 10mM Sr^2+^. In cases where [Ca^2+^]_i_ oscillations were induced by injection of mPLC*ζ* cRNA, eggs were placed in TL-Hepes media containing 2 mM Ca^2+^ within 20 minutes of the injection of mPLC*ζ* which occurred 4 hours post-injection of the FRET constructs (Camui or FRESCA).

### Pharmacological tests in mouse eggs

Mouse eggs were transferred to Ca^2+^ free TL-Hepes containing desired concetnrations of pharmacological compounds 5 min prior to Ca^2+^ imaging. FRET (YFP/CFP) was monitored simultaneously with Ca^2+^ (rhodamine signal). First, we determined how much inhibitor could be added without affecting Ca^2+^ entry. Concentrations of inhibitors were chosen based on this information as well as what was used in previous studies, see text for references. All media was Ca^2+^ free with the exception of PMA. The following concentrations were used: KN93 (0.5 μM), GO6983 (3 μM), Bim1 (5 μM), PMA (1 μM), AS105 (5 μM), AS461 (5 μM). AS105 has not been used in mouse eggs, so we adjusted the concentration to a level where the Ca^2+^ entry was not affected. Eggs with the first 4 compounds added were stimulated with 0.5 μM ionomycin 5 min after monitoring started, while the eggs with AS compounds were stimulated with 2.5 μM ionomycin. Side by side controls were performed under the same conditions (0.5 μM vs. 2.5 μM ionomycin).

### Data processing & statistical analyses

Graphs reporting FRET changes and Ca^2+^ responses were prepared using the values of the YFP (436×535)/CFP (436×480) ratios on the left axis, whereas Rhod-2 values were calculated using the following formula (F)/F0 (actual value at x time/average baseline values for the first 2 minutes of monitoring) and the scale placed on the right axis. Values from three or more experiments performed on different batches of eggs are presented as means ± s.e.m and were analyzed by the Student’s t-test. Differences were considered significant at P <0.05.

**Table 1.** Replicates in Camui experiments

| Camui | Experiment type | # of replicates | # of mice | # of total eggs |
| --- | --- | --- | --- | --- |
| | Iono 0.5 $\mu\text{M}$ | 3x | 2 | 12 |
| | Iono 2.5 $\mu\text{M}$ | 4x | 3 | 16 |
| | Iono 5.0 $\mu\text{M}$ | 3x | 2 | 12 |
| | $\text{Sr}^{2+}$ | 2x | 3 | 11 |
| | PLCζ | 3x | 3 | 16 |

**Table 2.** Replicates in FRESCA experiments

| **FRESKA** | **Experiment type** | **# of replicates** | **# of mice** | **# of total eggs** |
| --- | --- | --- | --- | --- |
| | Iono 0.5 $\mu\text{M}$ | 4x | 2 | 16 |
| | Iono 2.5 $\mu\text{M}$ | 4x | 3 | 16 |
| | Iono 5.0 $\mu\text{M}$ | 3x | 2 | 8 |
| | $\text{Sr}^{2+}$ | 3x | 3 | 22 |
| | PLCζ | 2x | 3 | 9 |
| Added compounds |  |  |  |  |
| Control | Iono 0.5 $\mu\text{M}$ | 5x | 2 | 27 |
| | Iono 2.5 $\mu\text{M}$ | 2x | 2 | 13 |
|  | PMA | 2x | 2 | 10 |
|  | BIM1 | 1x | 2 | 5 |
|  | KN93 | 2x | 2 | 10 |
|  | GO6983 | 2x | 2 | 9 |
|  | AS105 | 2x | 2 | 10 |
|  | AS461 | 2x | 2 | 15 |

## Supplement Figures

**Figure 2 – figure supplement 1.**
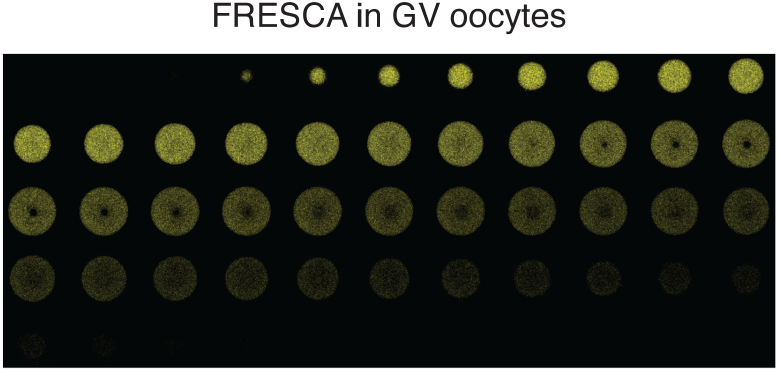
FRESCA expression in GV oocytes. Expression is mostly cytoplasmic, as well as some limited nuclear expression.

**Figure 2 – figure supplement 2.**
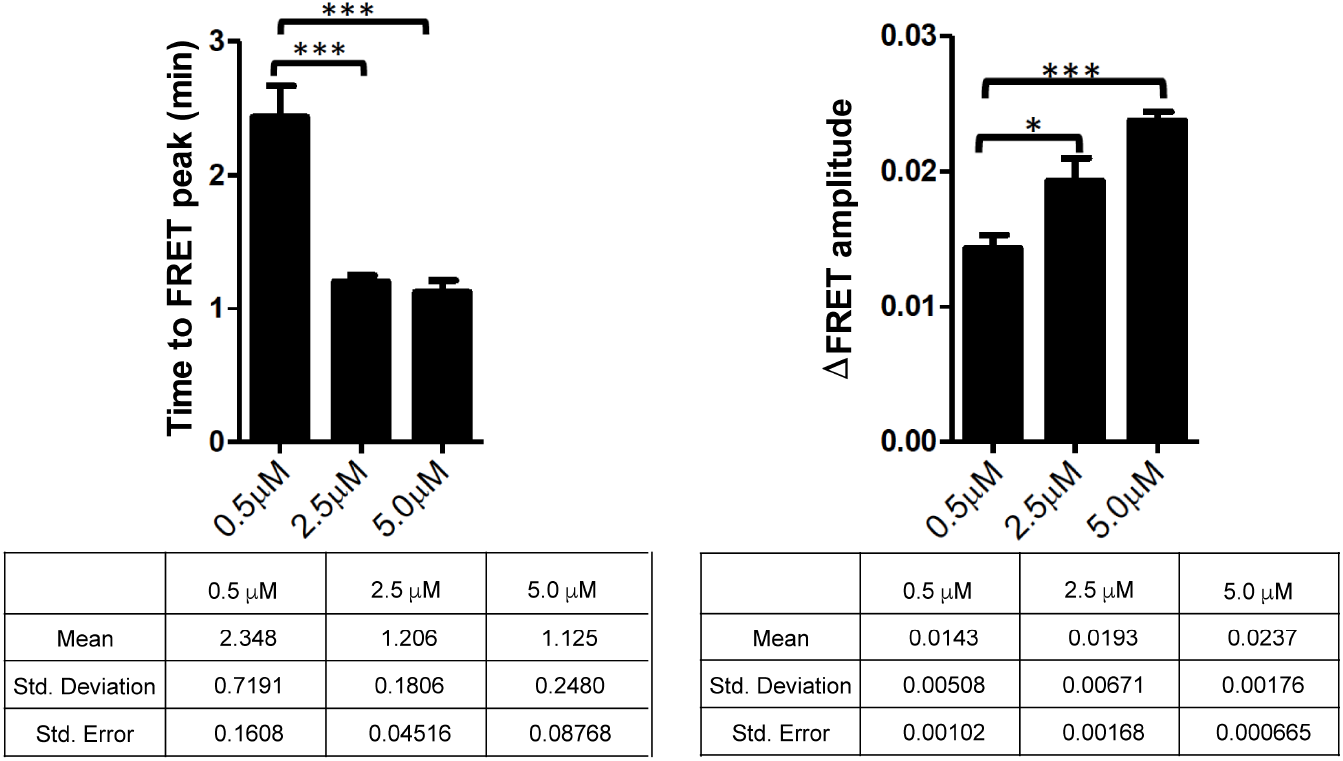
Quantification of FRESCA response to ionomycin addition. Time to FRET peak indicates how long it takes FRESCA to reach maximum ΔFRET signal after addition of ionomycin and increase in Ca^2+^. ΔFRET amplitude indicates the overall change in FRET during the duration of the Ca^2+^ signal. Statistics are reported in the tables below, differences were considered significant at P <0.05 (*).

**Figure 2 – figure supplement 3.**
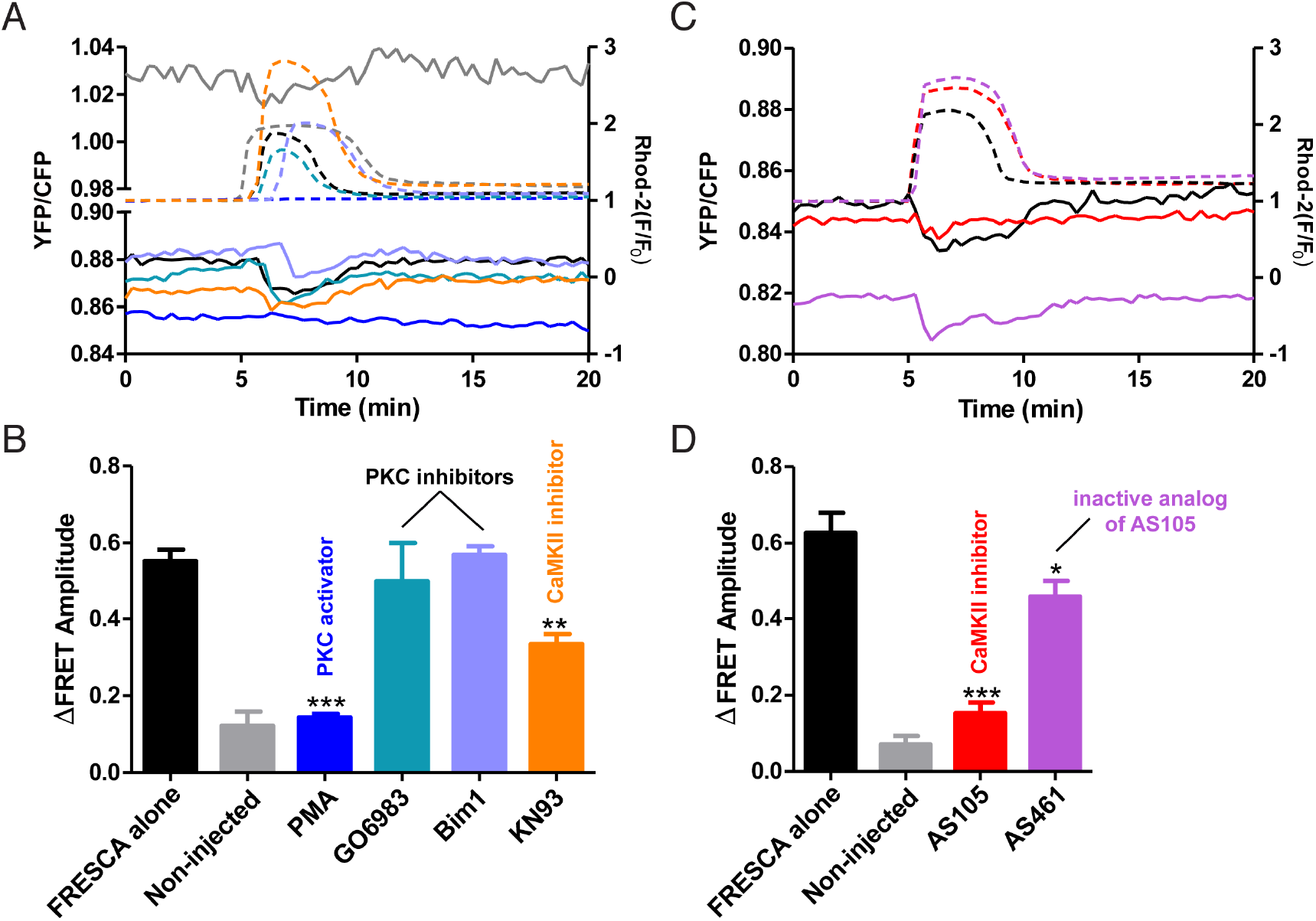
Pharmacological tests of FRESCA specificity in mouse eggs. Various compounds were added to mouse eggs expressing FRESCA. Ca^2+^ was monitored using Rhod-2 and FRET was monitored by YFP/CFP ratio. A) CaMKII inhibitor: KN93 (0.5 μM), PKC inhibitors: GO6983 (3 μM) and Bim1 (5 μM), and a PKC activator: PMA (1 μM) were added to mouse eggs and stimulated with 0.5 μM ionomycin. These were directly compared to FRESCA alone and a non-injected control with 0.5 μM ionomycin. Colors correspond to the bar graph in (B). B) Quantification of the amplitude of FRET signal in (A). C) CaMKII inhibitor: AS105 (5 μM) and inactive analog of this inhibitor: AS461 (5 μM) were added to mouse eggs and stimulated with 2.5 μM ionomycin. These were directly compared to FRESCA alone and a non-injected control with 2.5 μM ionomycin. Colors correspond to the bar graph in (D). D) Quantification of the amplitude of FRET signal in (C). Differences were considered significant at P <0.05 (*).

**Figure 3 – figure supplement 1.**
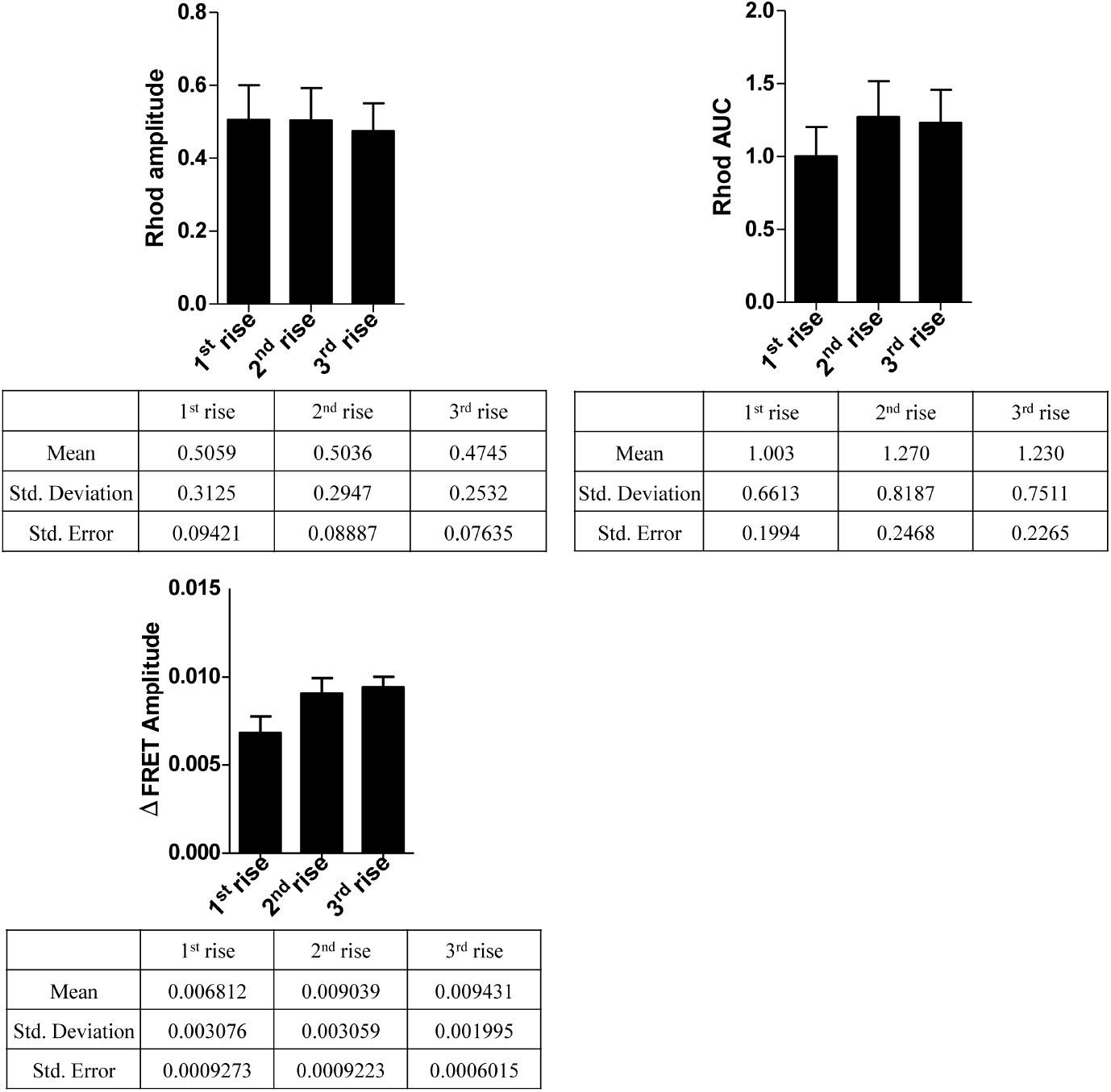
Quantification of Rhod-2 and FRET signals for FRESCA during Sr^2+^ induced oscillations. Three Ca^2+^ rises were quantified. The “1^st^ rise” is that which induced the first FRET response, and then the subsequent 2 rises were measured. Statistics are reported in the tables below, differences were considered significant at P <0.05 (*).

**Figure 3 – figure supplement 2.**
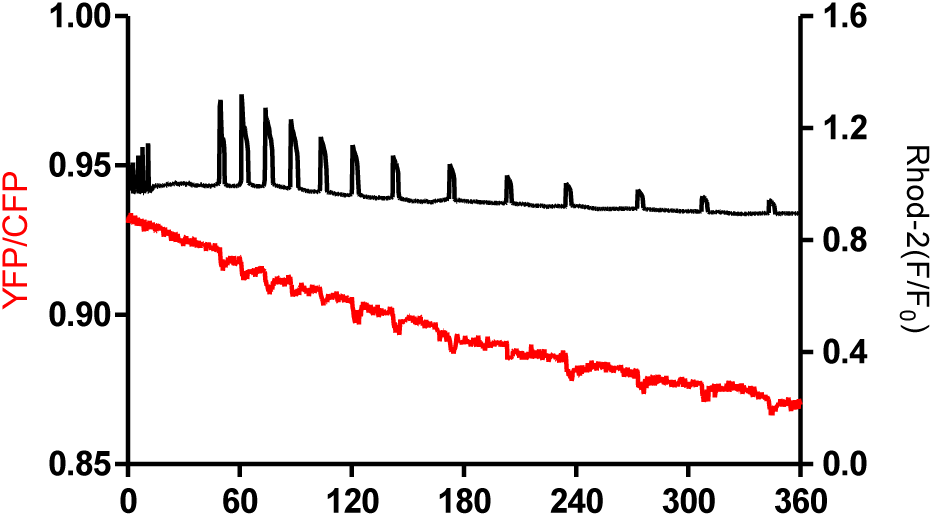
Increased time course of FRESCA response to Sr^2+^. Sr^2+^ (10 mM) was added to mouse eggs expressing FRESCA. Ca^2+^ is monitored by Rhod-2 (black line) and endogenous CaMKII activity is tracked by FRESCA (red line) for 6 hours. The progressive decreasing amplitude of the Rhod-2 AM fluorescence signal (especially after 180 min) is an artifact due to compartmentalization of the dye due to prolonged monitoring at >35 °C.

**Figure 4 – figure supplement 1.**
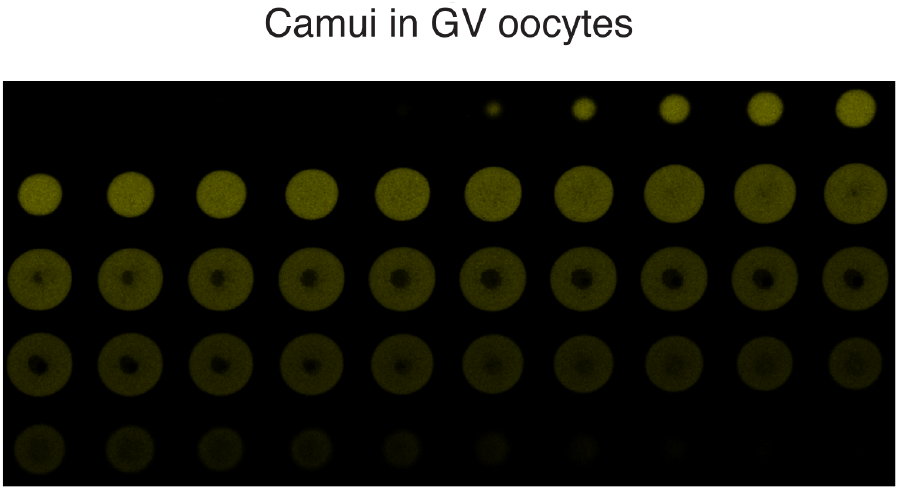
Camui expression in GV oocytes. Expression is mostly cytoplasmic, and is excluded from the nucleus.

**Figure 4 – figure supplement 2.**
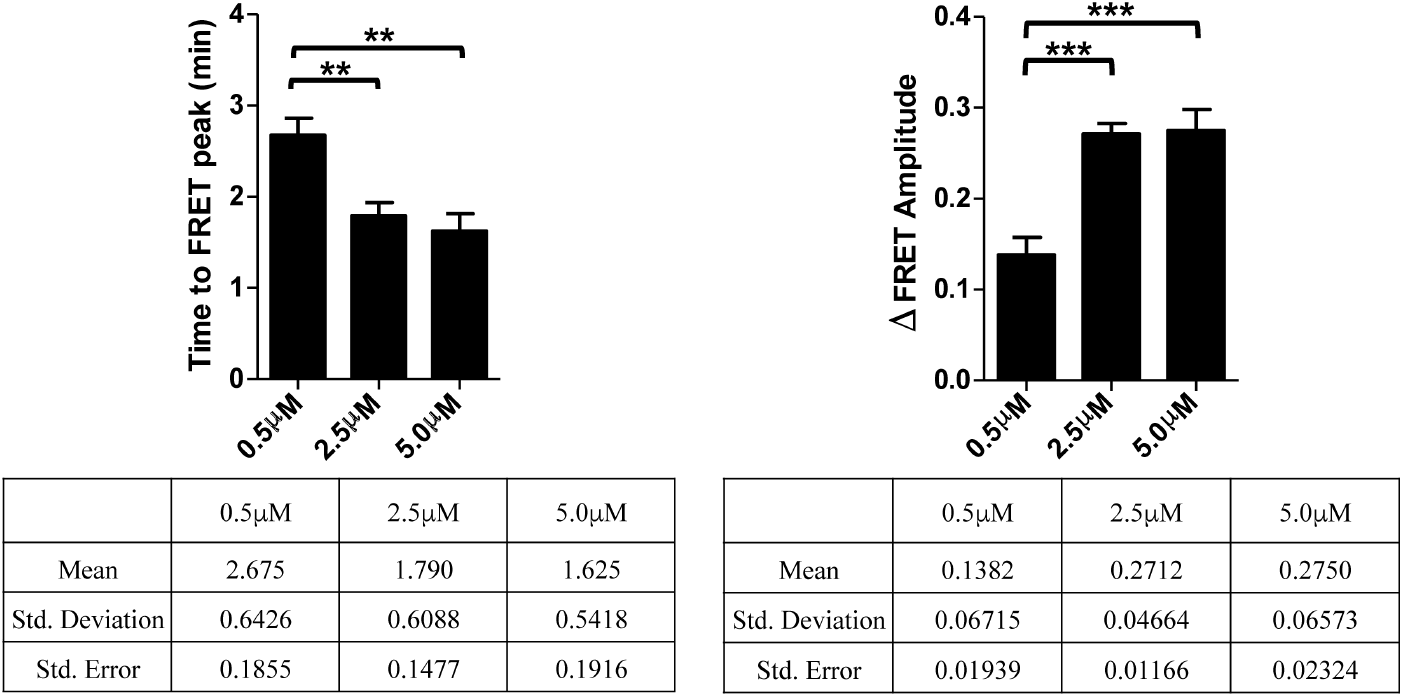
Quantification of Camui response to ionomycin addition. Time to FRET peak indicates how long it takes FRESCA to reach maximum ΔFRET signal after addition of ionomycin and increase in Ca^2+^. ΔFRET amplitude indicates the overall change in FRET during the duration of the Ca^2+^ signal. Statistics are reported in the tables below, differences were considered significant at P <0.05 (*).

**Figure 5 – figure supplement 1.**
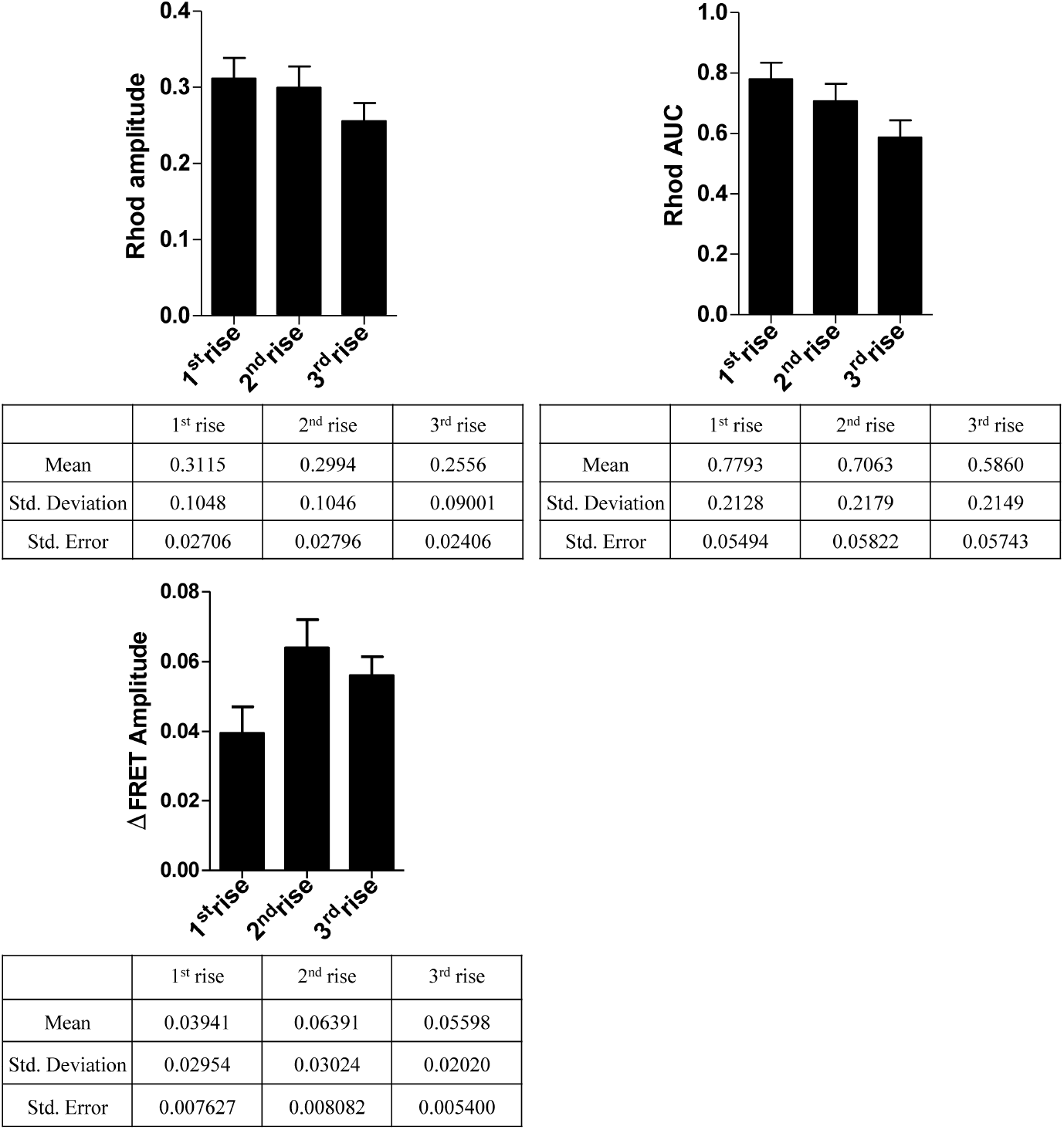
Quantification of Rhod-2 and FRET signals for Camui during Sr^2+^ induced oscillations. Three Ca^2+^ rises were quantified. The “1^st^ rise” is that which induced the first FRET response, and then the subsequent 2 rises were measured. Statistics are reported in the tables below, differences were considered significant at P <0.05 (*).

**Figure 6 – figure supplement 1.**
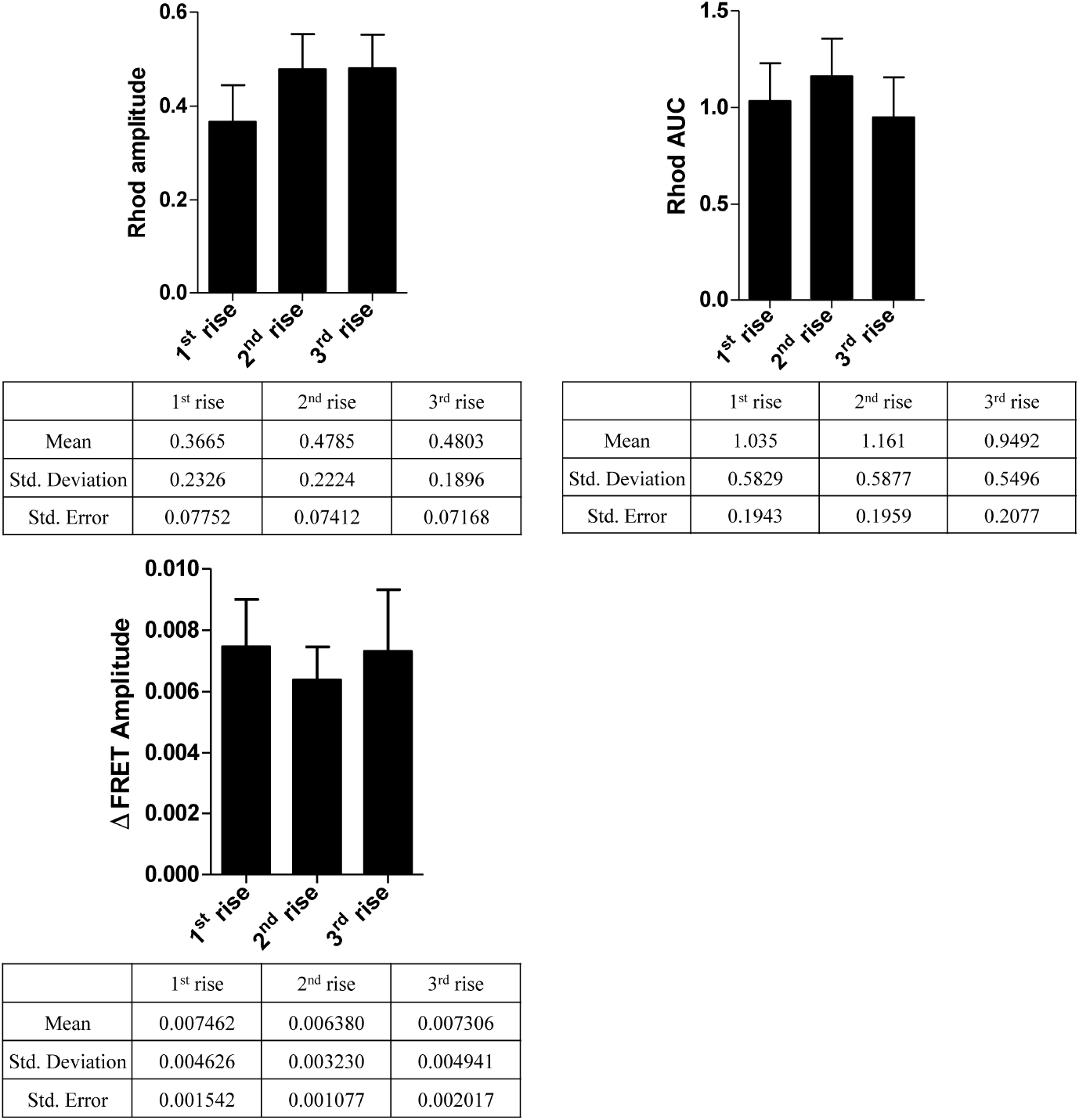
Quantification of Rhod-2 and FRET signals for FRESCA during PLC*ζ* induced Ca^2+^ rises. Three Ca^2+^ rises were quantified. The “1^st^ rise” is that which induced the first FRET response, and then the subsequent 2 rises were measured.

**Figure 6 – figure supplement 2.**
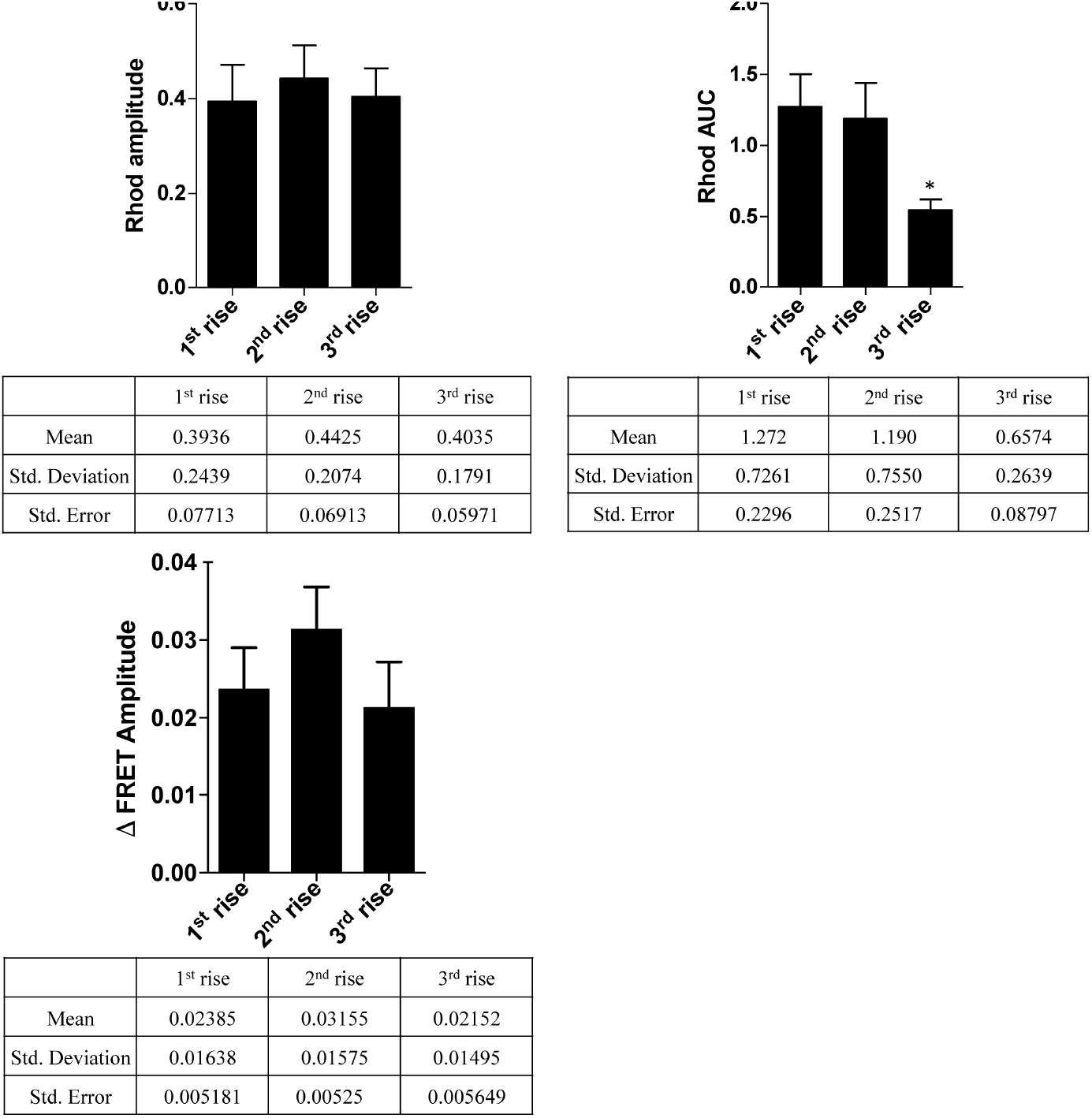
Quantification of Rhod-2 and FRET signals for Camui during PLC*ζ* induced Ca^2+^ rises. Three Ca^2+^ rises were quantified. The “1^st^ rise” is that which induced the first FRET response, and then the subsequent 2 rises were measured. Differences were considered significant at P <0.05 (*).

